# Spatial analysis of gut microbiome reveals a distinct ecological niche associated with the mucus layer

**DOI:** 10.1101/675918

**Authors:** Kellyanne Duncan, Kelly Carey-Ewend, Shipra Vaishnava

**Affiliations:** Molecular Microbiology and Immunology, Brown University, Providence, RI, 02912

## Abstract

Mucus associated bacterial communities are critical for determining disease pathology and promoting colonization resistance. Yet the key ecological properties of mucus resident communities remain poorly defined. Using an approach that combines *in situ* hybridization, laser microdissection and 16s rRNA sequencing of spatially distinct regions of the mouse gut lumen, we discovered that a dense microbial community resembling a biofilm is embedded in the mucus layer. The mucus associated biofilm-like community excluded bacteria belonging to phylum Proteobacteria. Additionally, it was significantly more diverse and consisted of bacterial species that were unique to it. By employing germ-free mice deficient in T and B lymphocytes we found that formation of biofilm-like structure was independent of adaptive immunity. Instead the integrity of biofilm-like community depended on Gram-positive commensals such as Clostridia. Additionally, biofilm-like community in the mucus lost fewer Clostridia and showed smaller bloom of Proteobacteria compared to the lumen upon antibiotic treatment. When subjected to time restricted feeding biofilm like structure significantly enhanced in size and showed enrichment of Clostridia. Taken together our work discloses that mucus associated biofilm-like community represent a specialized community that is structurally and compositionally distinct that excludes aerobic bacteria while enriching for anaerobic bacteria such as Clostridia, exhibits enhanced stability to antibiotic treatment and that can be modulated by dietary changes.

## Introduction

The past two decades have been a golden age for microbiome research. By comparing microbial composition between different physiologic states in humans and mouse models, it has become clear that various aspects of host physiology, metabolism, and immune system are intimately linked to the microbes and their interactions with the host (1). Accordingly, the gut microbiome has been shown to play a role in many western diseases, including inflammatory bowel diseases (IBD) such as Crohn’s and ulcerative colitis, colon cancer and metabolic diseases such as diabetes and obesity (2). However, attempts to identify a common pattern of microbial dysbiosis linked with these diseases have failed. Multiple studies show that bacterial communities in the gut are spatially organized and disrupted spatial organization of the gut microbiome is often a common underlying feature of disease pathogenesis (3–5). As a result, focus over the last few years has shifted from cataloging the diversity of gut bacteria towards understanding gut microbiome in spatial context (6). This line of investigation has brought the role of the intestinal mucus and associated microbial communities into sharp focus. A study involving the largest cohort of pediatric Crohn’s disease patients demonstrated that assessing the mucus-associated microbiome compared to fecal communities is more effective in early diagnosis of disease (7). Moreover, isolation of colonic mucus from mice mono-colonized with specific commensals showed that bacterial species present in the mucus show differential proliferation and resource utilization compared with the same species in the intestinal lumen of mice (8). In both mouse and human, it was shown that mucus-associated communities were distinct and not represented by fecal communities (9). Studies show that mucosal communities are refractory to colonization by probiotics unless perturbed by antibiotics (10). Although these studies underscore the critical role of mucus residing communities in determining disease pathology and promoting colonization resistance, ecological features of mucus resident communities that confer these properties remain poorly defined.

To fill this gap in our understanding of the gut microbiome ecology, we developed a novel strategy that combines fluorescence in-situ hybridization (FISH) with the spatial dissection power of laser capture microdissection (LCM) and 16S microbiome sequencing. We found that the microbial community closest to the host overlying the inner mucus layer forms a unique structure that harbors significantly higher species richness, selectively excludes bacteria belonging to class Gammaproteobacteria and whose integrity is dependent on anaerobic bacteria such as Clostridia. We propose that this community represents a previously unrecognized ecological safe harbor that promotes stability of the gut microbiome.

## Results

### Bacteria overlying the colonic mucus forms dense community structure that is compositionally distinct from adjoining community

Fluorescence *in-situ* hybridization (FISH) to 16s rRNA gene in combination with specialized tissue fixation methodologies that preserve mucus structure in an intact state enable the observation of microbe-microbe and host-microbe relationships *in-situ* (4, 11, 12). To understand the landscape of bacterial communities residing within the gut, we evaluated the spatial organization of gut bacteria in the transverse sections of mouse colon using universal 16s rRNA FISH probes (Figure 1A). FISH analysis of transverse sections of Carnoy’s fixed mouse colon showed that bacteria mixed in with digesta occupied the luminal space uniformly. However, a dense band of bacterial cells localized adjacent to the mucosa all around the tissue cross section. Quantitative measurement revealed that the dense band of bacteria extended up to 29 μm into the lumen (Figure 1A and B). The dense bacterial community resided at the boundary of inner and outer mucus layer (Figure 1C), and was on average 17μm thick in mice from our colony at Brown University (Figure 1D). Similar structure comprising of dense bacterial community close to the mucosa has been observed by FISH in mice by Swidsinski et al. which they named “interlaced layer” due to the bacterial organization being interlaced between the epithelium and fecal material (13). Furthermore, in the colon of gnotobiotic mice colonized with a defined 15-member community, bacteria were found to be concentrated at the border between the lumen and mucosa, (14), thus indicating that higher concentration of bacteria at the intersection of mucus layer and lumen is a conserved feature of microbial community organization within the mouse colon. In addition to being found within the mouse colon, such a structure has also been identified in rats, baboons, and humans (15) suggesting that mucus associated dense bacteria community is present throughout mammalian gut microbiomes. However, the ecological characteristics and importance of this community to the host have yet to be defined.

**Figure 1.**
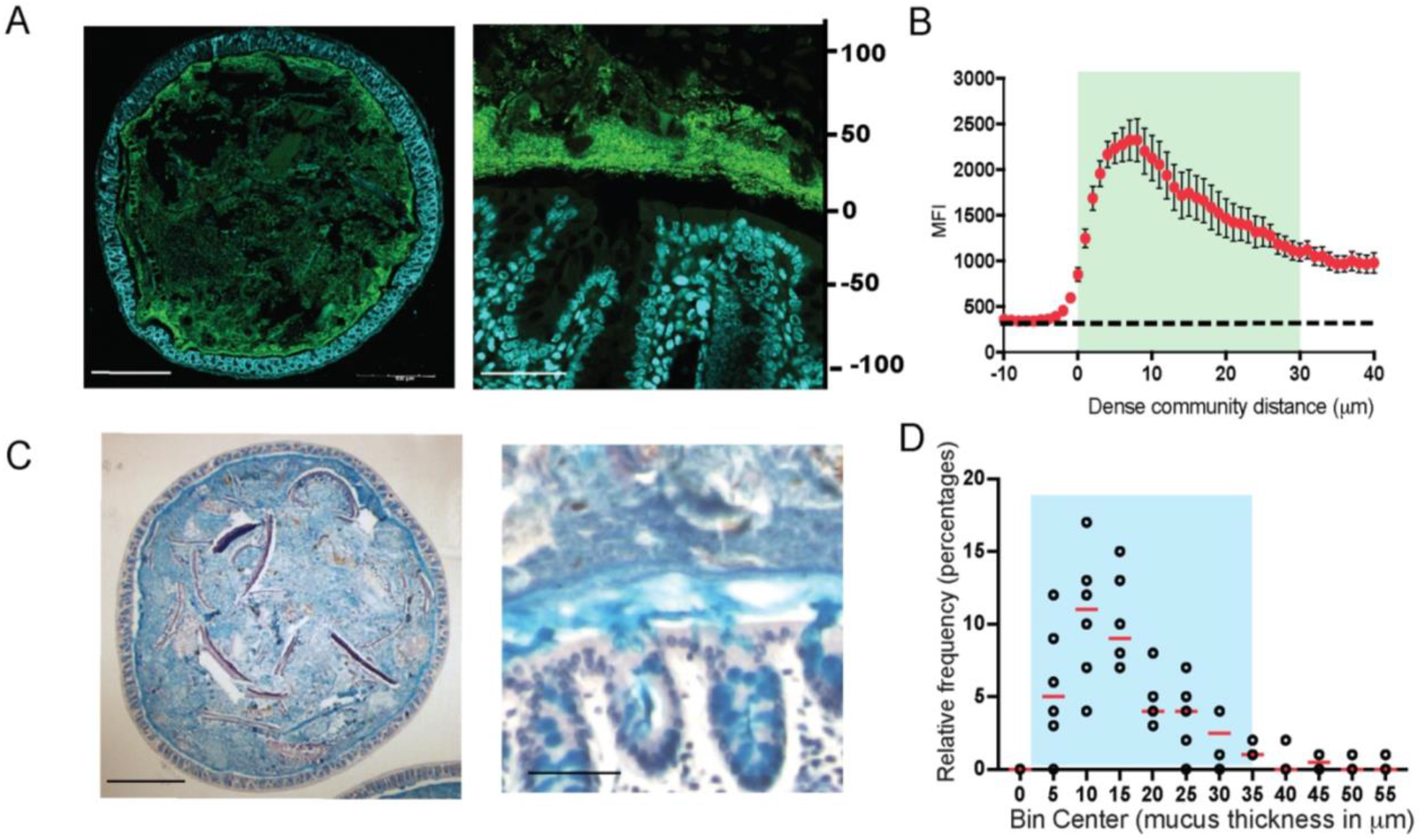
Dense band of bacteria forms overlying colonic mucosa. A. Colonic cross section stained with fluorescent probes identifying all bacteria (green) and host epithelium (blue) to show dense community structure close to the host (Left: 10x; Right: 60x). B. Mean fluorescence intensity (MFI) measurements from a representative mouse (average N=3) to show bacterial concentration of the dense community. Measurements start 10-μm before start of bacterial signal closest to epithelial lining. C. Colonic cross section stained with Alcian Blue/Pas to show dense mucus layer (Left: 10x; Right: 60x). D. Relative frequency of dense mucus layer thickness measurements, described in Methods section under “Mucus Thickness” (N = 6 mice, 40 measurements each).

To assess the microbial composition of the dense bacterial community we adapted laser-capture microdissection (LCM) technique to isolate defined regions of the colonic mucus layer followed by 16S rRNA sequencing. The LCM isolation strategy was designed so as to capture the first 100μm from the epithelial lining corresponding to the observed thickness of the dense bacterial community abutting inner mucus layer (inner) independently of the adjoining 50μm section of sparse community colonizing the outer mucus layer (outer) (Figure 2A-B). Using this strategy, we collected an area of 1,000,000 μm^2^ of the inner community and corresponding 500,000-μm^2^ of the outer community for each mouse colon. qPCR quantification of the community samples confirmed that the inner community had higher bacterial load compared to the outer community (Figure 2C). We applied similar dissection strategy to colonic tissue sections from germ-free (GF) mice to ensure that the LCM procedure did not introduce any bacterial DNA contamination (Supp. Fig. 1). We isolated the inner and outer communities from mice from two different cages and compared microbiome dissimilarities based on cage and location (inner vs outer). We used the distance metric UniFrac that uses phylogenetic relationships to evaluate differences in overall composition between communities (16). Using weighted UniFrac distances that calculates dissimilarities in abundance of microbes (quantitative), we only saw a difference in community composition between the two cages but no difference was noted between the two locations (Figure 2D). The unweighted analysis that calculates dissimilarity based on presence or absence of a microbe (qualitative) showed that location was the second most contributing variable that significantly separated the inner community from the outer community (Axis 2) (Figure 2E). Evaluating composition without abundance indicates differences in communities is based on which taxa are able to live in them (17). Differential abundance analysis at class level revealed that the inner dense community just like the outer sparse community comprised mainly of bacteria belonging to class Bacteroidia and Clostridia. However, the outer sparse community harbored significantly more Gammaproteobacteria compared to the inner dense community (Figure 2F and G, Supp. Fig. 1B, C). These results suggest that Gammaproteobacteria are selectively restricted from colonizing the mucus layer closest to the host epithelium.

**Figure 2.**
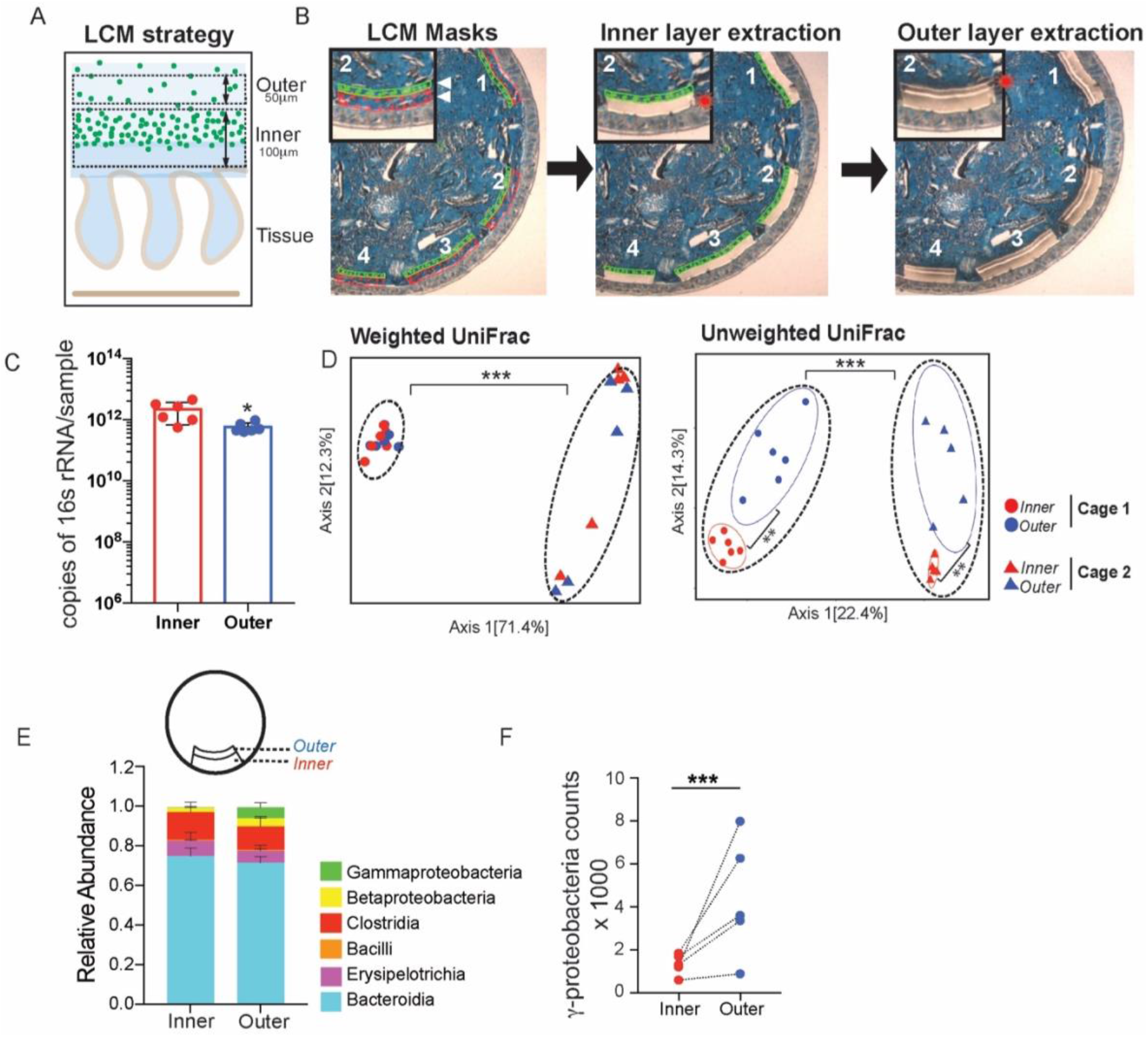
Mucus associated dense community is compositionally distinct from adjoining sparse community. A. Diagram of laser capture microdissection (LCM) strategy for independently obtaining the inner dense community (1^st^ 100-μm) and outer sparse community (subsequent 50-μm) of the mucosal layer for 16S rRNA sequencing. B. Images of colonic cross sections showing LCM masks of inner and outer communities followed by the independent extraction of each region separately in order to analyze regions separately. C. Copy number of 16S rRNA per sample from inner and outer communities showing enough material is obtained for sequencing, with more copies in the inner community (N=6 mice). *p < 0.05 paired t-test. D. Principal coordinates of analysis (PCoA) plots. Left: Weighted UniFrac shows most significant difference in microbial communities comes from cage effect, with no difference in location. Right: Unweighted UniFrac shows most significant difference in microbial communities comes from cage effect, but also the inner and outer communities are significantly different (N = 11 mice). **p < 0.01, ***p < 0.001 PERMANOVA. E. Class level relative abundance of inner and outer communities (N = 5 mice from Cage 2). F. Differential abundance of class Gammaproteobacteria calculated using DESeq2, showing significantly more Gammaproteobacteria in the outer community (N = 5 mice from Cage 2, Log2 Fold Change = 4.87, padj = 1.39 x 10^-7^). ***p < 0.001.

### Dense bacterial community closest to the host has significantly higher species richness due to presence of low abundance unique amplicon sequence variants (ASVs)

Diversity is one of the most important ecological attributes which determines stability of a community (18). The number of species represented in an ecosystem (its richness) and the way in which individuals are distributed amongst the species (its evenness) are often combined into a single index to quantify diversity. Plotting the number of unique amplicon sequence variants (ASVs) found at increasing subsampling depths with an alpha rarefaction curve showed that the inner community consistently had more unique observed ASVs (Figure 3A). The Chao1 index estimates the whole population community richness, we saw the inner community that corresponds with the dense bacterial structure has significantly higher Chao 1 index (Figure 3B, Supp. Fig. 1D). Species evenness in a community captures another aspect of diversity by determining diversity as a standardized index of relative species abundance. We evaluated the evenness of the two communities by applying Pielou’s evenness index (Pielou, 1969) and found the inner community is significantly less even than the outer community (Figure 3D, Supp. Fig. 1F). Accordingly, Shannon-Weaver, a commonly used diversity index that considers both richness and evenness of communities, showed no difference between the two communities. (Figure 3C, Supp. Fig. 1E).

**Figure 3.**
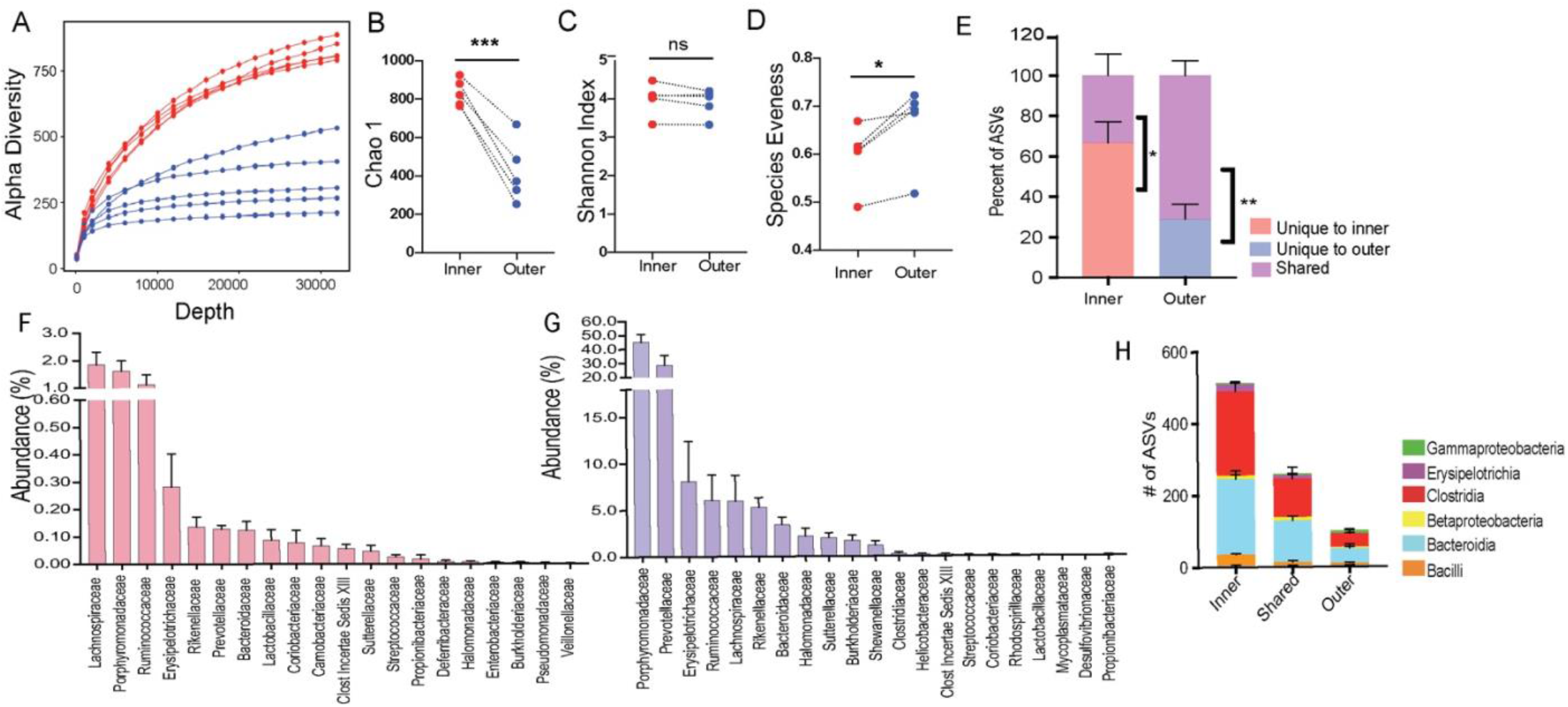
Inner dense community has significantly higher species richness due to presence of low abundance unique amplicon sequence variants (ASVs). A. Alpha rarefaction curves plotting the number of unique amplicon sequence variants (ASVs) from inner (red) and outer (blue) communities, showing more unique ASVs found in the inner community (N = 5 mice from Cage 2). B. Chao1 diversity index showing significantly higher richness in the inner community (N = 5 mice from Cage 2). ***p < 0.001 paired t-test. C. Shannon diversity index showing no difference in Shannon diversity between inner and outer communities (N = 5 mice from Cage 2). D. Pielou’s Evenness index showing significantly higher evenness in the outer community (N = 5 mice from Cage 2). *p < 0.05 paired t-test. E. Proportion of ASVs found either unique to either region or shared within each mouse showing the inner community maintains a large proportion of unique ASVs and the outer community mostly consists of ASVs also found in the inner (N = 5 mice from Cage 2). *p < 0.05, **p < 0.01 paired t-test. F. Abundance of unique ASVs in the inner community of each mouse condensed by family to show the unique ASVs of the inner community are at low abundance (N = 5 mice from Cage 2). G. Abundance of shared ASVs between the inner and outer communities of each mouse condensed by family to show the shared ASVs comprise the majority of the community (N = 5 mice from Cage 2). H. Number of ASVs found either unique to either region or shared within each mouse condensed by class showing the increased richness in the inner community is comprised of unique ASVs from classes Clostridia and Bacteroidia (N=5 mice from Cage 2).

Next, we wanted to determine the number of ASVs that were unique or shared between the inner and outer community within each mouse. We found in each animal significantly more ASVs were unique to the inner dense (average 70% unique, 30% shared) (Figure 3E) whereas the adjoining sparse region comprised of more shared ASVs (30% unique and 70% shared). Members of classes Bacteroidia and Clostridia comprise the majority of bacteria that are either unique to one location or shared between both communities (Figure 3F). Additionally, at family level unique microbes made up a very small percent (<0.2%) of the total abundance within the inner dense community (Figure 3G). Conversely, microbes shared between the two locations were highly abundant within the dense inner community (Figure 3H). It could be that many microbial species in the outer sparse community are too scarce to be captured by sequencing and that we detect these low abundant microbial species only when their relative abundance increases in the inner dense community. Therefore, to ensure that the increased diversity found in the inner community was not artificially inflated due to the larger biomass captured, we used mice from Jackson Laboratories that have much thinner inner mucus layer mucus layer (~5μm) but still harbor a comparable dense community at the intersection of inner and outer mucus layer (Supp. Fig. 2 A-C). We isolated 50μm area from the epithelium corresponding to inner community corresponding and the adjacent 50μm for as outer community. We saw that even when there was no significant difference in the biomass of the two regions when amplifying the 16S rRNA gene (Supp. Fig. 2 D) the inner community was still significantly more diverse than the outer community (Supp. Fig. 2E-G). Thus, our results demonstrate that mucus associated dense community represent a novel species bank of hidden microbial diversity and functional potential in the gut.

### Formation of biofilm-like community structure is independent of adaptive immunity

To evaluate the role of adaptive immunity in the formation of the dense structure, we used Rag KO mice, containing a disruption in the recombination activating gene 1 (Rag1), therefore unable to initiate V(D)J rearrangement of immunoglobulin and T-cell receptors and fail to generate mature B-cells or T-cells. (19). Proinflammatory T-cells (Th1, Th17) contribute to pathogen clearance by orchestrating an immune response, but need to be regulated by T-regulatory (Tregs) to limit excessive inflammation (20). By suppressing the immune response, Tregs facilitate the colonization of commensals (21) and preserve diversity within the gut microbiome (22). Immunoglobulin A (IgA) secreted from B-cells enables pathogen clearance (23, 24) but also facilitates colonization and persistence of commensals (25).

The immune system of a germ-free animal is considered naïve due to the lack of education usually provided by the microbiota (26). The adaptive immune response takes at least 4-7 days to be fully activated (27), so we evaluated Rag KO mice after 4-weeks of conventionalization (Figure 4A). After 4 weeks, Rag KO mice form a comparable biofilm-like structure close to the host epithelium to that of wild-type mice, showing that B-cells and T-cells do not influence the formation of the biofilm-like structure (Figure 4B-C). Comparing the fecal microbiome communities recovered, we found the composition was significantly different when evaluating with weighted metric but not when using the unweighted metric (Supp. Fig. 3A-B), suggesting the differences in genotype influence the abundances but not colonization of the microbiota. The most notable difference at family level was in the abundance of Prevotellaceae which previously been identified as a bacteria that are highly coated by secretory IgA (28, 29). To understand the role of adaptive immunity in shaping the composition of the mucosal community, we used LCM to isolate 100 μm from the host epithelium. We found that like the fecal communities, the mucosal community showed a significant difference when using the weighted metric but not the unweighted metric (Figure 4D-E). Similar to fecal communities the family responsible for the divergence in the mucosal layer were Prevotellaceae (Figure 4F-G). Both mucosal and fecal communities are influenced by the adaptive immune system in regards to the abundance of microbes but not their presence. Additionally, the ability of the dense community structure to form in both innate and adaptive immune deficient mice suggests this assembly is independent of adaptive component such as secretory IgA.

**Figure 4.**
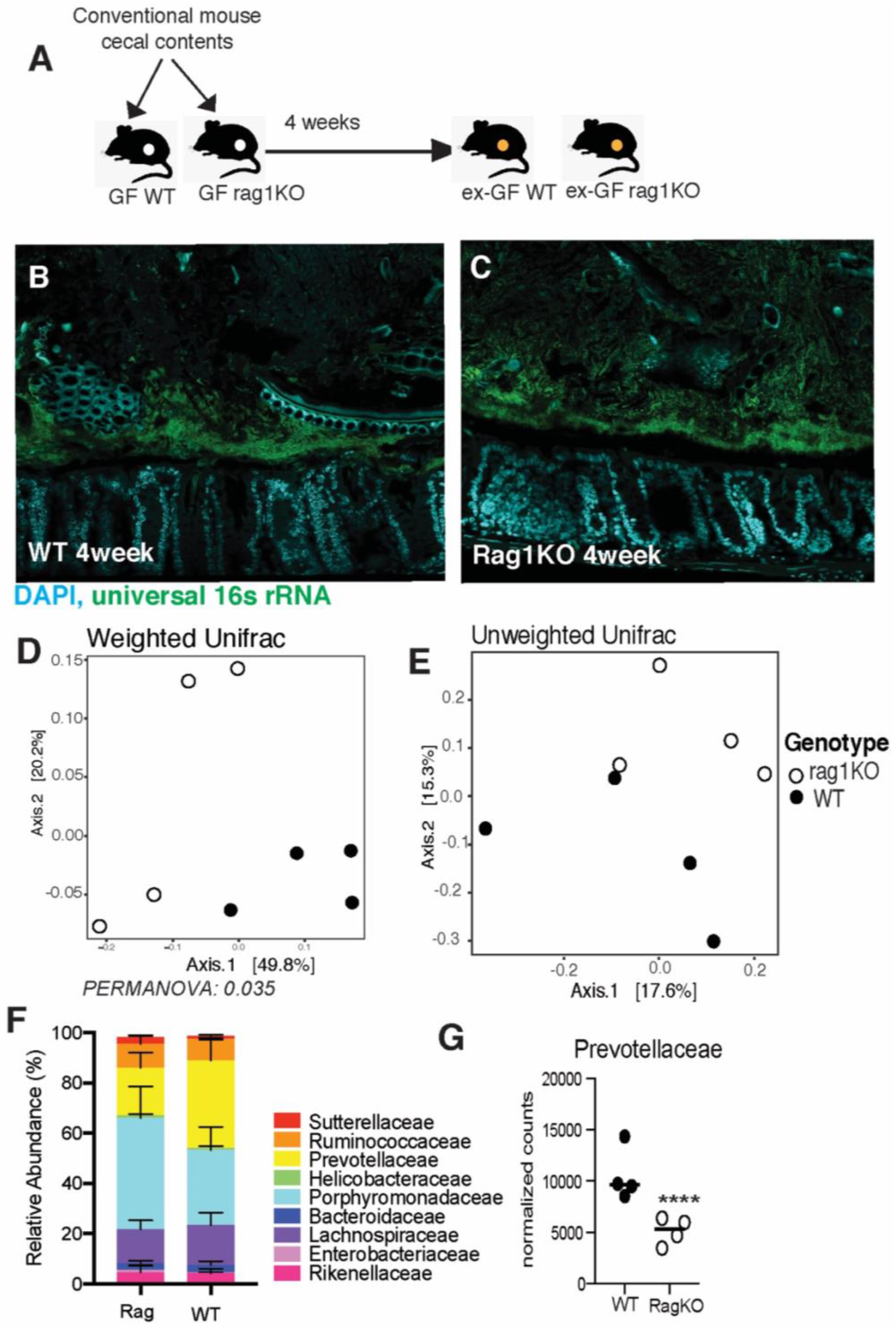
Mice deficient in adaptive immunity still form biofilm-like community with some changes to composition. A. Experimental design for comparing dense community formation in WT and Rag1KO mice. Provided germ-free WT and Rag1KO mice with same donor flora and examined community structure after 4 weeks of conventionalization. B. Colonic cross section stained with fluorescent probes identifying all bacteria (green) and host epithelium (blue) to show dense community structure forms after 4 weeks in WT conventionalized mice. C. Colonic cross section stained with fluorescent probes identifying all bacteria (green) and host epithelium (blue) to show dense community structure forms after 4 weeks in Rag1KO conventionalized mice. D. Principal coordinates of analysis (PCoA) plot using weighted UniFrac distances shows there is a significant difference between mucosal communities of conventionalized WT and Rag1KO mice (N = 4 mice per group). *p < 0.05 PERMANOVA. E. Principal coordinates of analysis (PCoA) plot using unweighted UniFrac distances shows there is no significant difference between mucosal communities of conventionalized WT and Rag1KO mice (N = 4 mice per group). p = 0.113 PERMANOVA F. Family level relative abundance of mucosal communities in conventionalized Rag1KO and WT mice (N = 4 mice per group). G. Differential abundance of family Prevotellaceae calculated using DESeq2, showing significantly less Prevotellaceae in the mucosal communities of Rag1KO mice (N = 4 mice per group mice, Log2 Fold Change = 1.0, padj = 6.67 x 10^-5^).

### Clostridia are important for maintaining structural integrity of biofilm-like community

In order to assess the contribution of community composition in the maintenance of the dense band of bacteria, we subjected it to perturbation by a broad (Ciprofloxacin) and narrow (Vancomycin) spectrum antibiotic (Fig. 5A). Ciprofloxacin disrupts both DNA gyrase and topoisomerase IV function, therefore disrupting replication in all microbes making it a broad spectrum antibiotic (30). Vancomycin disrupts peptidoglycan synthesis, most prominent in Grampositive organisms like Clostridia and therefore is a narrow-spectrum antibiotic (31). Comparison of fecal communities of mice treated with Vancomycin or Ciprofloxacin to that of untreated communities confirmed that both antibiotics significantly decreased the diversity of the microbiome, a characteristic of antibiotic treatment (Fig. 5B-C). The mice treated with Vancomycin experienced a larger disruption in fecal community composition compared to those treated with Ciprofloxacin (Fig. 5C). Vancomycin treatment as expected resulted in a significant depletion of Clostridia and an expansion of Gammaproteobacteria (Fig. 5C). To determine if differences in community composition between Ciprofloxacin and Vancomycin treatments correlated with differences in community structure, we employed FISH to visualize the gut microbiome *in-situ*. Ciprofloxacin treated communities maintained the dense bacterial community structure similar to that seen in untreated mice (Figures 5D), whereas this structure was completely lost in mice treated with Vancomycin (Figures 5E). The Ciprofloxacin-treated mice were devoid of Proteobacteria (Figures 5D, panel ii) whereas the remaining community in Vancomycin-treated mice harbored members of Proteobacteria (Figures 5E, panel ii). Additionally, Clostridia were preserved within the dense community upon Ciprofloxacin treatment (Figures 5D, panel iv), but lost after Vancomycin treatment (Figures 5E, panel iv). Obliteration of the dense community structure is correlated with the loss of Clostridia within the dense community. These data suggest that vancomycin-sensitive microbes such as Clostridia are important for maintaining community structure.

**Figure 5.**
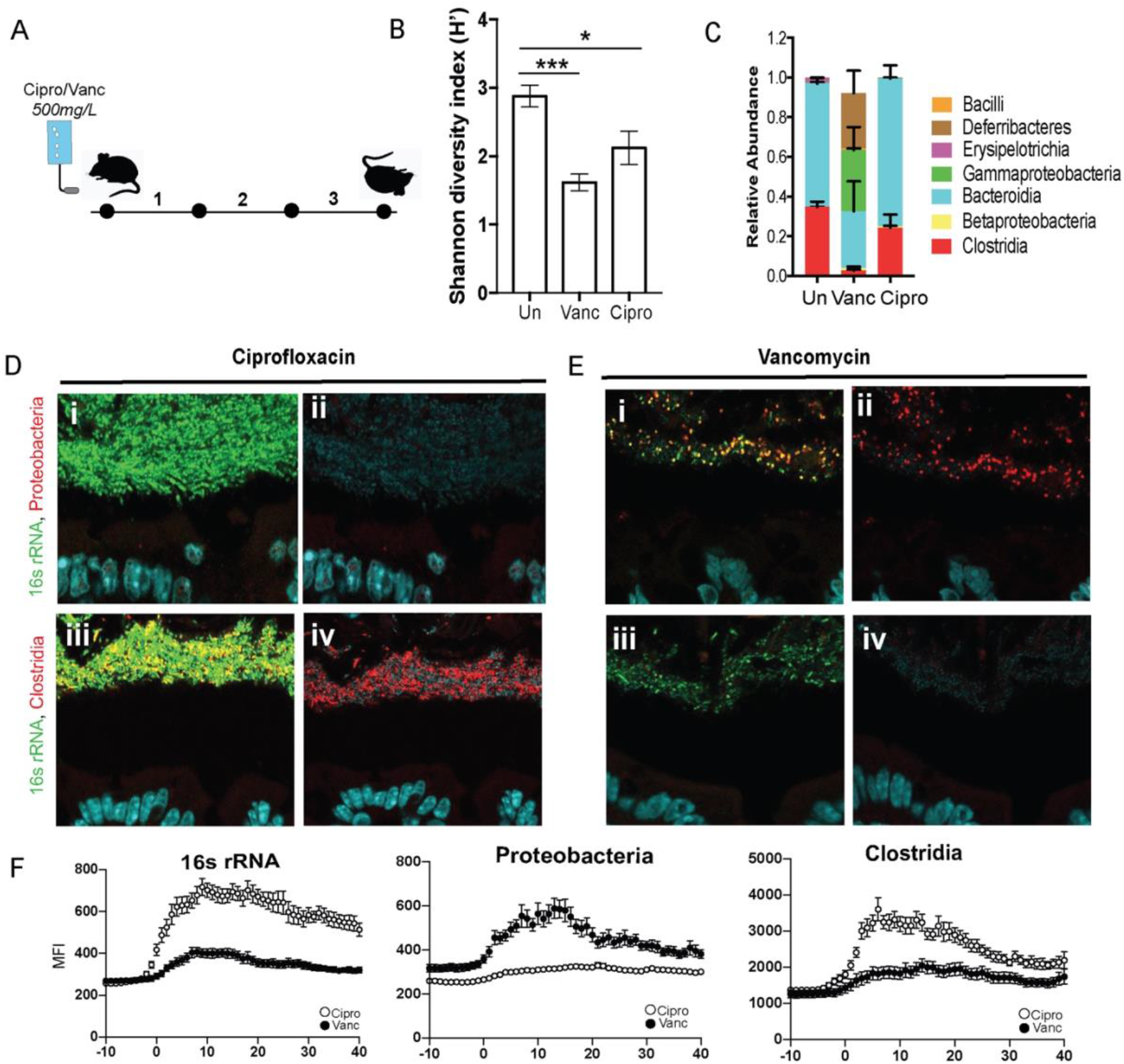
Dense community structure is sensitive to Vancomycin. A. Experimental design for comparing dense community structure in Ciprofloxacin and Vancomycin treated mice. Provided SPF WT mice with 500mg/L Ciprofloxacin or Vancomycin in the drinking water for three days. B. Shannon diversity of untreated, Vancomycin, and Ciprofloxacin treated fecal communities (N = 6 mice per group). C. Class-level relative abundance for untreated, Vancomycin, and Ciprofloxacin treated fecal communities (N =6 mice per group). D. FISH images of spatial structure after Ciprofloxacin treatment showing preservation of community consisting of Clostridia and excluding Proteobacteria. *i:* merged image with host epithelium (DAPI, blue), all bacteria (16s rRNA, green), and Proteobacteria (red). *ii*. Merged image with host epithelium and Proteobacteria. *iii*. Merged image with host epithelium, all bacteria, and Clostridia (red). *iv*. Merged image with host epithelium and Clostridia E. FISH images of spatial structure after Vancomycin treatment showing decimation of dense band of bacteria as well as the absence of Clostridia and the prevalence of Proteobacteria. *i*. merged image with host epithelium (DAPI, blue), all bacteria (16S rRNA, green), and Proteobacteria (red). *ii*: merged image with host epithelium and Proteobacteria. *iii*. merged image with host epithelium, all bacteria, and Clostridia (red). *iv*. Merged image with host epithelium and Clostridia F. Mean fluorescence intensity (MFI) measurements for representative mice quantifying bacterial density for all bacteria (16s rRNA), Proteobacteria, and Clostridia after Ciprofloxacin (open circles) and Vancomycin (closed circles) treatment. Measurements started 10-μm before start of bacterial signal closest to epithelial lining (N = 3 mice).

### Mucus associated dense community maintains higher species richness and is less prone to dysbiosis than luminal community following antibiotics

Uneven distribution or spatial heterogeneity of species within an ecosystem increases its stability during large scale perturbations (32). We wanted to assess whether the spatial heterogeneity we observe within the gut mucosal community contributes to its stability. Mice from Jackson laboratories were subjected to a clinically relevant vancomycin treatment for 10 days (33)(Figure 6A). Considering the thinner mucus layer in Jackson mice, (Figure S2), we isolated the mucus associated community by capturing 50μm section closest to the intestinal epithelium. Luminal community was isolated by excising a 300μm diameter circle from the center of the cross section. Antibiotic treatment significantly altered the mucus community as well luminal communities (Figure 6B-C). Before antibiotic treatment communities in both locations had similar richness, however following antibiotic treatment the mucus associated community lost significantly less richness than the luminal communities (Figure 6D). The greater impact of antibiotic treatment on species richness in the luminal community suggests the dense community is less susceptible to antibiotic perturbation. Additionally, after antibiotic treatment, we saw a significantly increase in the relative abundance of Proteobacteria and a decrease in the relative abundance of Firmicutes in the luminal community compared to the mucus associated dense community (Figure 6E). Specifically, we saw that post antibiotic treatment, the differential abundance of Clostridia was significantly more in the mucus associated dense community than the lumen community (Figure 6F). In contrast, a larger bloom β-Proteobacteria was observed in the lumen compared to the mucus associated dense community (Figure 6G). Quantification with qPCR and Clostridia specific 16s rRNA FISH further confirmed that after antibiotic treatment significantly more Clostridia are preserved within the mucus compared to lumen (Figure 6H-I). Our results thus reveal that mucus associated microbial community suffers less disruption than the luminal microbial community.

**Figure 6.**
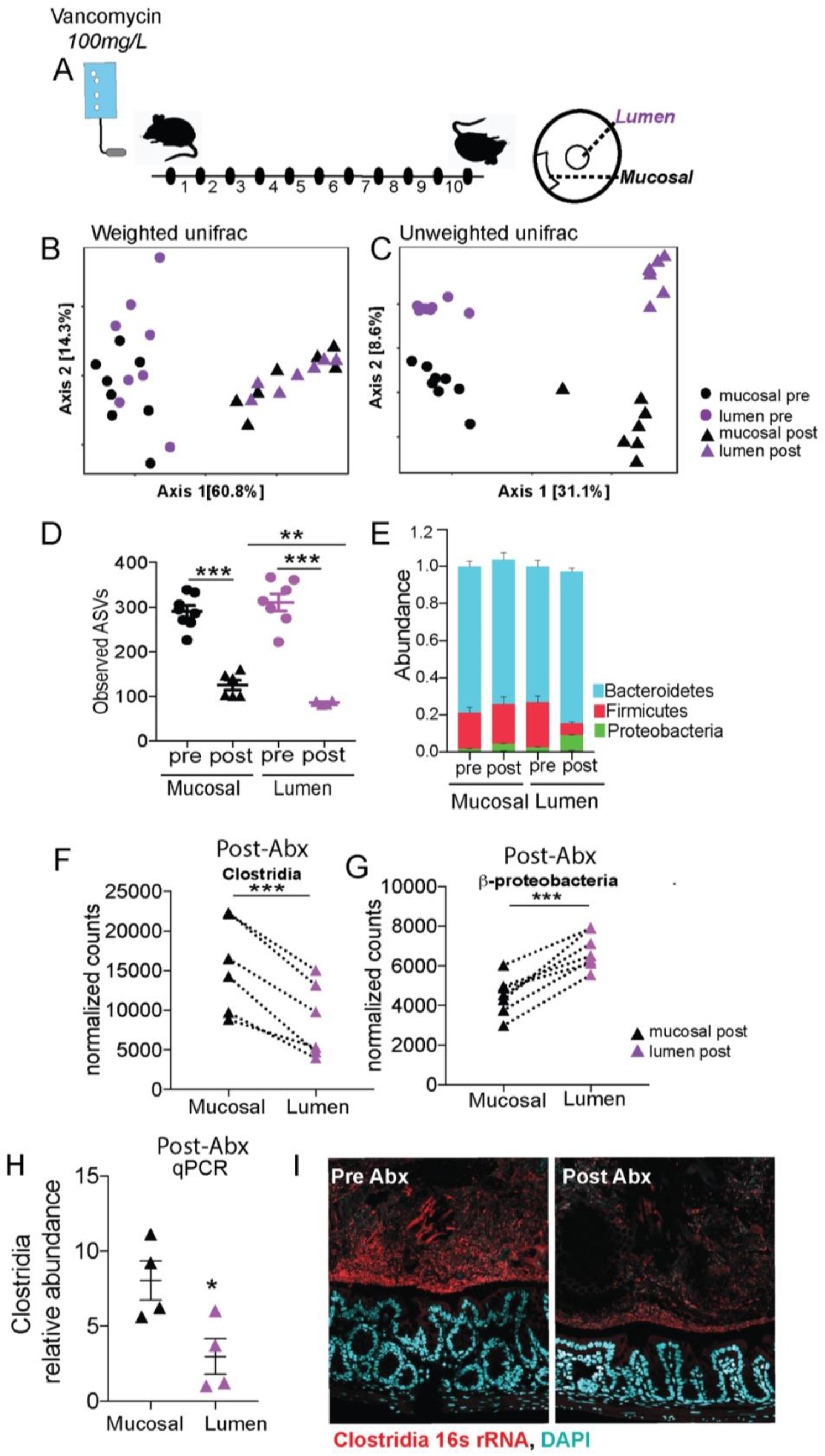
Following antibiotic treatment inner dense community maintains higher species richness and is less prone to dysbiosis than luminal community. A. Experimental design for comparing effects of antibiotics on spatially distinct regions of the gut microbiome. Provided mice water with 100mg/L of Vancomycin in drinking water. Sacrificed after 10 days of treatment. B. Principal coordinates of analysis (PCoA) plot using weighted UniFrac distances shows both mucosal and luminal communities are significantly altered by antibiotic treatment (N = 7 mice per group). C. Principal coordinates of analysis (PCoA) plot using unweighted UniFrac distances shows both mucosal and luminal communities are significantly altered by antibiotic treatment (N = 7 mice per group). D. Number of unique ASVs found in each sample at rarefied depth of 39,464 reads (N = 7 mice per group) showing a significant depletion in richness in both locations following antibiotic treatment (paired t-test) and the luminal community lost significantly more ASVs than the mucosal community following antibiotic treatment (unpaired t-test). **p < 0.01, *** p < 0.001. E. Phylum-level relative abundance for pre- and post-antibiotic mucosal and luminal communities (N = 7 mice per group). F. Differential abundance of class Clostridia, showing significantly more Clostridia in the mucosal community after antibiotic treatment (N = 7 mice per group, Log2 Fold Change = 1.33, padj = 0.034). ***p < 0.001 DESeq2. G. Differential abundance of class β-Proteobacteria, showing significantly more β-Proteobacteria in the luminal community after antibiotic treatment (N = 7 mice per group, Log2 Fold Change = 0.59, padj = 0.034). ***p < 0.001 DESeq2. H. qPCR quantification of Clostridia relative abundance of mucosal and luminal communities after antibiotics, showing increased Clostridia in the mucosal community following treatment (N = 4 mice per group). *p < 0.05 paired t-test. I. FISH images of mucosal communities before and after Vancomycin treatment showing the greater prevalence of Clostridia (red) before treatment and the reduction after antibiotics, and the greater maintenance of Clostridia in the dense community adjacent to the host epithelium (DAPI, blue) following treatment.

### Time restricted feeding structurally enhances biofilm-like community and enriches for Clostridia

To create a model where the dense community structure could be manipulated, we investigated dietary interventions that would potentially increase the thickness of the dense community structure without modifying the nutrient proportions as to limit changes to the microbiome composition due to variations in microbial metabolism. When microbial populations are well-fed, they grow more rapidly and thus lose diversity and spatial structure (34). Additionally, reduced nutrient availability can establish nascent biofilms (35). To exploit these known characteristics of spatial structure in microbial communities, we used intermittent fasting (IF) to determine whether this dietary intervention would increase the thickness of the dense structure compared to *ad libitum* (AL) feeding.

Mice were subjected to 16:8 IF (16 hours fasting, 8 hours feeding) every day for 30 days (Figure 7A). Mice are nocturnal and do the majority of their feeding at night, so the 8 hours of feeding was scheduled during the dark cycle (36). As a control, cage-mates were maintained in a separate cage with constant access to food for *ad libitum* feeding. Bedding was switched daily between the two cages to minimize drift in microbiome composition. The amount of food eaten was measured daily for each group, and we found there was no difference in the amount of food eaten between the IF and AL mice (Figure 7B). Mice were weighed daily after IF mice finished feeding and we found no significant difference between the body weights of the IF and AL mice (Figure 7C). There is a trend for IF mice to have increased body weights, however this may be an artifact of the increased food consumption of IF mice during their feeding period since the IF mice eat the same amount as the AL mice but restricted to the 8-hour feeding period. Intermittent fasting is known to increase insulin sensitivity and therefore reduces blood glucose levels (37). To determine whether 4-weeks was a sufficient amount of time to see this metabolic phenotype, we performed a glucose tolerance test on both groups of mice to examine their ability to absorb glucose. We found at peak levels of glucose in the blood (30 minutes), AL mice had significantly higher blood glucose levels than the IF mice (Figure 7D), suggesting the 4 weeks of IF was sufficient for inducing glucose sensitivity.

**Figure 7.**
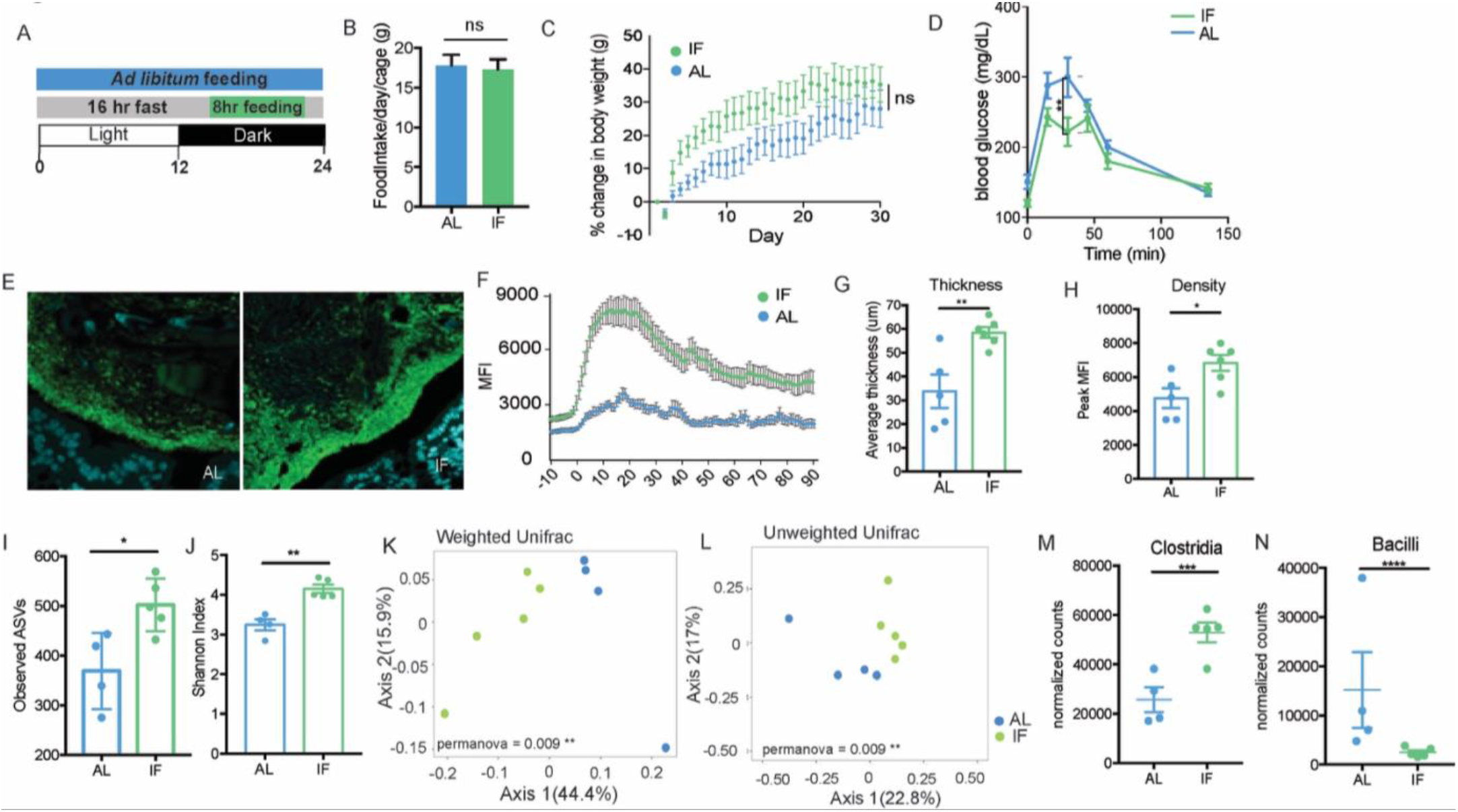
Intermittent fasting increases thickness and density of biofilm-like community and significantly alters community composition. A. Experimental design for intermittent fasting experiment. Intermittent fasting (IF) mice were allowed *ad libitum* feeding during an 8-hour feeding window during the dark cycle and fasted for the remaining 16 hours. This was repeated every day for 30 days. A control group was given continuous *ad libitum* (AL) feeding. B. The number of grams of food eaten per 24-hour period by each cage showing no significant difference in the amount consumed between AL and IF mice. C. Percent change in body weight over the course of the experiment showing no significant difference in the increase in body weight between AL and IF mice. D. Blood glucose levels taken before, 15min, 30min, 45min, 60min, and 130min post glucose injection showing a significant increase in blood glucose levels of AL mice at 30min post-injection, suggesting reduced insulin sensitivity compared to IF mice (N = 6 mice per group). *p < 0.05, **p < 0.01 2-way ANOVA. E. Colonic cross section stained with fluorescent probes identifying all bacteria (green) and host epithelium (blue) of dense community structure with Left: *Ad libitum* feeding and Right: Intermittent fasting, showing increase in thickness and density with IF treatment. F. Overlay of MFI measurements from representative IF and AL mice showing difference in thickness and density. G. Thickness of dense community structure after AL and IF measured by MFI showing significantly thicker communities after IF (N = 5-6 mice per group). *p < 0.05 unpaired t-test. H. Peak MFI measurements of dense community structure after AL and IF showing significantly denser communities after IF (N = 5-6 mice per group). *p < 0.05 unpaired t-test. I. Observed ASVs found in AL and IF LCM-captured communities (150-μm) showing significantly higher richness in IF communities. Samples were rarefied to lowest sampling depth for normalization (N = 4-5 mice per group). *p < 0.05 unpaired t-test. J. Shannon diversity in AL and IF LCM-captured communities (150-μm) showing significantly higher diversity in IF communities. Samples were rarefied to lowest sampling depth for normalization (N = 4-5 mice per group). **p < 0.01 unpaired t-test. J. Principal coordinates of analysis (PCoA) plot using weighted UniFrac distances shows LCM-captured communities (150-μm) from AL and IF are significantly different (N = 4-5 mice per group). **p < 0.01 K. Principal coordinates of analysis (PCoA) plot using unweighted UniFrac distances shows LCM-captured communities (150-μm) from AL and IF are significantly different (N = 4-5 mice per group). **p < 0.01 L. Differential abundance of Clostridia between LCM-captured communities (150-μm) from AL and IF communities, showing significantly more Clostridia in IF communities (N = 4-5 mice per group, Log2 Fold Change = 1.04, padj = 0.0017). ***p < 0.001 DESeq2. K. Differential abundance of Bacilli between LCM-captured communities (150-μm) from AL and IF communities, showing significantly less Bacilli in IF communities (N = 4-5 mice per group, Log2 Fold Change = 2.59, padj = 0.0004). ****p < 0.0001 DESeq2.

To determine whether IF increased the thickness of the biofilm-like community, we used FISH to visualize and quantify the size of the community structure close to the epithelial lining. We found that four weeks of IF significantly increased the thickness and density of the dense community structure (Figure 7E-H, S4 A-B). The increased dense community structure following intermittent fasting provides a model for evaluating the role of the dense community structure within the gut microbiome.

Intermittent fasting has been shown to induce changes in microbiome composition, including increasing the species richness of the community (38). To understand how the microbiome changes in a spatial context with IF, we evaluated changes in both the biofilm-like structure and fecal communities. In fecal communities, there is no significant difference in diversity or composition between IF and AL communities when using a weighted distance, but is significantly different when using an unweighted metric, suggesting the communities differ based on the presence of low abundant microbes (Supp. Fig 4). To evaluate the composition of the biofilm-like community, we used LCM to isolate 150-μm from the epithelial lining and IF communities close to the host are significantly more diverse than AL communities (Figure 7I-J). LCM captured IF and AL communities are significantly distinct using both weighted and unweighted distance metrics (Figure 7K-L), where IF mice have significantly increased abundance of *Clostridia* and less *Bacilli* in the biofilm-like community than AL mice (Figure 7M-N). Together these data show the differences in community composition imposed by IF are more strongly affecting the biofilm-like structure close to the host opposed to fecal communities. The expansion of the biofilm-like structure coincides with the expansion of *Clostridia*, suggesting *Clostridia* expansion could be the mechanism for increasing the biofilm-like structure.

## Discussion

Epithelial surfaces in the gastrointestinal tract are covered by a layer of mucus, which prevents bacteria from accessing mucosal surface (11, 39). Mucins are chemically and structurally diverse molecules, mostly consisting of linear and branched oligosaccharides and are important source of carbohydrate for saccharolytic bacteria in the gut (40). Large complex polymers such as mucin first need to be degraded by several different hydrolytic enzymes to smaller oligomers, monosaccharides, and amino acids before they can be assimilated by intestinal microorganisms(41). In the gut breakdown of mucin is a result of cooperative activity of multiple members of the gut microbiota (42, 43). Studies using two-stage continuous fermentation system show that fecal bacteria rapidly adhere and colonize mucin gel forming mucin-degrading biofilms (43).

Biofilms constitute the dominant mode of microbial life in most ecosystems (44, 45) yet they remain poorly described in the gut that harbor dense consortium of commensal bacteria. We used a classical histological procedure to preserve mucus integrity, specific techniques for microbiota detection (i.e., FISH) and UV-laser assisted dissection of biofilm communities in micron scale for 16s rRNA microbiome analysis. Our method using entire cross sections of mouse colon fixed in Carnoy’s fixative to preserve mucus architecture allowed us to survey the structure and spatial composition of the gut microbiome as it exists *in-situ*. We saw that intestinal bacteria form a continuous biofilm-like structure, lining the mucus surface that coats the colonic mucosa. Our observation that mucus associated biofilm in the murine colon is a normal feature is in contrast to studies that associate intestinal biofilms with pathogenic phenotypes such as IBD and colon cancer (13, 46). However in these studies it was noted that the biofilms associated with diseased pathology had higher concentration of pathobionts such as *Bacteroides fragilis, E.coli* and *Fusobacterium nucleatum* compared to controls. Moreover, biofilms from patients with Crohn’s disease or with ulcerative colitis were shown to be disrupted, allowing pathobionts to spread and invade intestinal epithelia and cause inflammation (47). Thus, indicating that it is not the biofilm phenotype per se that drives the disease phenotype but structural and functional disruption of intestinal biofilms that drives disease pathogenesis in the gut. In fact, mucus associated biofilms provide an efficient strategy for commensals to resist environmental perturbations, the host immune system, and luminal flow in the GI tract and thus promote stability of the gut microbiota (48). Additionally, close proximity of mucus associated biofilms benefit the host by modulating immunity and acting as deterrent to colonization and invasion by enteropathogens (49).

Using antibiotics with different targets we found that the “narrow-spectrum” antibiotic induced the largest changes in biofilm structure and the loss of biofilm structure was correlated with ablation of Clostridia and the expansion of Proteobacteria. Biofilm-like structure was stable when treated with Ciprofloxacin, a broad-spectrum antibiotic, but collapsed completely when treated with Vancomycin that targets commensals belonging to class Clostridia. Collapse of biofilm structure coincided with bloom of Proteobacteria near the mucus. These results suggest that “narrow-spectrum” and “broad-spectrum” require new definitions that go beyond understanding how antibiotics specifically target certain organisms, and incorporate how they affect community architecture. A number of healthy people naturally carry *Clostridium difficile* in their large intestine and don’t have ill effects (50). *C.difficile* infection in humans are associated with recent antibiotic use (51). Once infected, *C.difficile* is known to form biofilms in the gut that resist antibiotic treatment (52). It would be important to understand whether the adverse effect of an antibiotic regimen on the commensal biofilm is directly correlated with susceptibility to *C.difficile* infection. Also equally important to determine would be whether fecal microbiota transfer (FMT) that corrects *C.difficile* infection restores homeostatic commensal biofilm.

16s rRNA microbiome analysis of the bacterial community micro-dissected from the biofilm revealed that Gammaproteobacteria, a bacterial class associated with antibiotic induced dysbiosis, are in much lower abundance in the biofilm compared to adjoining areas. In a healthy gut, members of phylum Proteobacteria are normally maintained at very low levels but expand significantly in dysbiosis (53, 54). Increased abundance of class Gammaproteobacteria is seen in metabolic disorders (55, 56), obesity (57, 58), and inflammatory bowel disease (7), all of which are pathologies characterized by intestinal inflammation (57). Our results suggest that biofilm structure rich in anaerobic commensals such as Clostridia suppress Proteobacteria bloom. Depletion of Clostridia within the biofilm is coincident with Proteobacteria bloom. We also found that antibiotic induced depletion of species richness and associated dysbiosis was significantly lower in biofilm compared to the lumen communities. Several studies show that biofilm-grown cells have properties that are distinct from planktonic cells, one of which is an increased resistance to antimicrobial agents (59). Work on *in-vitro* biofilms show that slow growth and/or induction of an rpoS-mediated stress response could contribute to antimicrobial resistance (60). Moreover, the physical and/or chemical structure of mucus where the biofilms are embedded could also confer resistance by exclusion of antibiotics from the bacterial community. Finally, bacteria growing as biofilm might develop a specific antibiotic-resistant phenotype (61). It is likely that there are multiple resistance mechanisms at work that confer this property to mucus biofilms and future work looking at meta-transcriptional response of mucus associated biofilms in health and disease would be crucial in further delineating these properties.

Mechanisms regulating the biofilm formation of commensal bacteria are yet to be fully elucidated. Whether these mechanisms are bacteria or/and host intrinsic remains to be seen. Bacterial intrinsic mechanisms such as expression of Serine-Rich Repeat Proteins (SRRP’s) that bind to host epithelial proteins in pH-dependent manner can be foreseen in helping biofilm formation in different niches of the gut (62, 63). Another bacterial intrinsic mechanism that could be crucial for biofilm formation is a mode of biochemical communication between different bacteria species called quorum sensing. The role of quorum sensing has been well established in forming biofilms in several bacterial species, including *E. coli, P. aeruginosa*, and *Salmonella sp*. (64). Quorum sensing between different species of gut commensals was recently shown to influence the abundance of the major phyla of the gut microbiota (65). On the host side, several innate and adaptive immune effectors that are continuously secreted into the gut lumen could play a role in entrenching the biofilm in the intestinal mucus. Recently it was shown that *in vivo* coating with IgA promotes aggregation of commensal bacteria *B.fragilis* within the mucus (25). A different study linked IgA binding to bacteria within the mucus layer of the colon to transcriptional changes that stabilize in microbial consortium (66). We saw that 4-weeks post colonization of WT and Rag KO germ-free mice with same donor fecal microbiome, comparable biofilm formed in the colon of two groups. Compared to WT mice, Rag KO mice showed reduction in relative abundance of bacteria belonging to family Prevotellaceae in both fecal as well as LCM extracted mucosal microbiomes. Bacteria belonging to Prevotellaceae were previously shown to be preferentially coated by IgA (28). Our results indicate that a comparable biofilm-like structure forms even when adaptive immunity is completely lacking. However, as previously reported we also find that absence of secretory IgA in the gut influences relative abundance of IgA coated bacterial members (67).

Gut epithelium also secretes a slew of antimicrobial proteins that are important for regulating spatial organization of bacteria that could play a critical role in biofilm formation. For example, RegIIIγ, a c-type lectin that specifically targets Gram positive bacteria regulates hostmicrobe spatial segregation in the small intestine could promote biofilm structure by maintaining immune tolerance of microbial communities close to the epithelium (4). A small lectin-like protein ZG16 (zymogen granulae protein 16) secreted by the host was recently shown to aggregate bacteria in the colonic mucus layer to maintain bacteria at a safe distance from the epithelial cell surface (68). Finally, epithelium secreted effector Ly6/PLAUR domain containing 8 (Lypd8) protein binds flagellated microbiota and keeps them away from epithelium (69). If and how components of innate immunity such as antimicrobial protein secreted by the intestinal epithelium effect mucosal community structure and composition of the biofilm-like structure remains to be determined.

Several studies have shown that in addition to the type of diet, the timing of food intake plays a critical role in shaping intestinal microbial ecology(70–72). We find that when mice were on time restricted feeding cycle the mucus associated biofilm-like structure significantly increases in size and intensity compared to mice on unrestricted access to food. This enhancement coincided with significant increase in Clostridia specifically in the biofilm-like structure suggesting that targeting Clostridia by dietary changes one could manipulate mucus associated biofilms. Time restricted feeding or intermittent fasting has been shown to alter the T cells in the gut resulting in reduction of IL-17 producing T cells and an increase in regulatory T cells and result in protection from experimental autoimmune encephalomyelitis (EAE) (73). It is tempting to speculate that increase in regulatory T cells could be due to increase in short chain fatty acid (SCFA) producing Clostridia in the mucus associated biofilm.

This study was done in mice, but the dense community structure has been observed in healthy humans (74). However, following the required preparation, biopsy washes of mucosal surfaces in healthy humans is basically sterile (75) and therefore makes it impossible with current technologies to determine whether these same ecological characteristics that we find in mice are also seen in humans. We recommend a reasonable solution would be to “humanize” germ-free mice with a human donor microbiota in order to determine whether human microbial communities are able to assemble in the same fashion.

These and other discoveries on the mechanisms orchestrating the gut microbiota spatial organization should be evaluated for their role in promoting homeostatic biofilms in the gut. Determining mechanisms that promote or disrupt homeostatic biofilm structure in gut should help towards identifying novel therapeutic targets in a broad variety of disorders mediated by microbiota biofilm dysbiosis.

## Methods

### Contact for Reagent and Resource Sharing

Further information and requests for resources and reagents should be directed to and will be fulfilled by the Lead Contact, Dr. Shipra Vaishnava.

### Experimental Model and Subject Details

#### Mice

All mice used were wild-type with a C57BL/6 background. Mice used for homeostatic characterization were bred in the SPF barrier facility at Brown University and sacrificed at 8 weeks of age. The mice from Cage 1 consisted of 6 male mice, all littermates and kept in the same cage. The other 5 are designated as Cage 2 and consisted of females from two different litters and kept in the same cage post-weaning (cage-mates). Mice used for characterization of spatial organization in Jackson mice were ordered from Jackson Laboratories, consisting of 3 males and 3 females and sacrificed at 6 weeks of age. Mice used for comparing mucosal and luminal communities after antibiotic treatment were ordered from Jackson Laboratories, consisting of 15 6-week old female cage-mates. Mice used for comparing Ciprofloxacin and Vancomycin antibiotic treatment were ordered from Taconic Biosciences, consisting of 12 7-week old female cagemates. Mice used for intermittent fasting experiments were ordered from Taconic Biosciences and started treatment when mice were 4-weeks old. Germ-free mice for conventionalization were bred in the Germ-free facility at Brown University ranging from 8-12 months old. Experiments were performed according to protocols approved by the Institutional Animal Care and Use Committees of Brown University.

#### Antibiotics Experiments

Experiment comparing Vancomycin and Ciprofloxacin antibiotic treatment was done by exchanging drinking water with either 500mg/L Vancomycin or 500mg/L Ciprofloxacin (pH 11).

Mice were sacrificed after 3 days of treatment. Experiment comparing mucosal and luminal communities after antibiotic treatment was done by exchanging drinking water with 100mg/L Vancomycin. Mice were sacrificed after 10 days of treatment.

#### Conventionalization Experiment

Experiment comparing community structure in conventionalized wild-type (WT) and Rag-1 KO (Rag1KO) mice was done by collecting cecal content from conventional mice and suspending in 8mL PBS before filtering through 70-um filter and gavaging 200-uL per mouse. Mice were sacrificed after 4-weeks post conventionalization

#### Intermittent Fasting Experiment

Experiment comparing intermittent fasting to *ad libitum* feeding was done by maintaining groups in separate cages and removing food from intermittent fasting mice at 5:30am and returning it at 9:30pm every day for 30 days. Bedding was switched every day between the intermittent fasting and *ad libitum* cages to minimize drift.

### Method Details

#### Sample Preparation

The distal colon was carefully removed to avoid disrupting architecture of encased contents and placed in 3mL Methacarn for 24 hours. Tissues were washed 3x over 3 days with 70% EtOH at 4°C. Tissues were processed at the Brown University Molecular Pathology Core with the Leica ASP300S by first dehydrating (70% EtOH 1 hr, 2 washes 95% EtOH 45 min, 1 hr, 3 washes 100% EtOH 45 min, 1 hr, 1 hr) then infiltrated with xylenes (3 washes, 45 min, 45 min, 1 hr) followed by paraffin wax at 60°C (1 hr, 1 hr, 1hr 15 min). Samples were embedded in paraffin to produce cross sections by holding tissues vertically in a base of hot wax while chilling. Once wax base secured the tissue vertically, the cassette was filled the rest of the way and chilled until hard.

#### Fluorescence in-situ hybridization and MFI Quantification

7-μm thick slides were deparaffinized through a solution series for 10 minutes each (xylenes, xylenes, 100% EtOH, 95% EtOH, 70% EtOH, H_2_O). Slides were placed in a pre-warmed (56°C) humidifying chamber and incubated 10 minutes with pre-warmed hybridization buffer (0.9M NaCl, 20mM Tris pH 7.4, 0.1% SDS). Hybridization buffer was removed and pre-warmed probe solutions were applied to regions of tissue isolated with hydrophobic slider marker. Probe solutions were prepared using 2μL EUB338I (1μg/μL), 2μL EUB338II (1μg/μL), 2 μL EUB338III (1μg/μL), and 94μL hybridization buffer per slide for viewing all bacteria (green) (76); 2μL EUB338I (1μg/μL), 2μL EUB338II (1μg/μL), 2 μL EUB338III (1μg/μL), 2μL GAM42a (1μg/μL), 2μL BET42a (1μg/μL), 35μL formamide, and 55μL hybridization buffer per slide for viewing all bacteria (green) and Proteobacteria (red) (77); 2μL EUB338-I (1μg/μL), 2μL EUB338-II (1μg/μL), 2 μL EUB338III (1μg/μL), 2μL Clep866 (1μg/μL), 2μL Erec482 (1μg/μL), 30μL formamide, and 60μL hybridization buffer per slide for viewing all bacteria (green) and Clostridia (red) (78, 79). Incubated slides at 56°C with probe solutions in humidifying chamber for 4 hours. Washed slides twice for 10 minutes in wash buffer (0.9M NaCl, 20mM Tris pH 7.4). Incubated slides with DAPI (1μg/mL) 10 minutes, followed by a final wash in wash buffer for 10 minutes. Dried slides 10 minutes at 56°C before applying coverslip with Fluro-Gel. Mean Fluorescence Intensity (MFI) measurements were taken using a Zeiss Inverted Microscope (Axio Observer Z1) using the profile function within the AxioVision Rel. 4.8 software to take the average MFI of the default 10-μm wide area, starting 10μm prior to the presence of bacteria and 40μm into the luminal space. 20 measurements were taken per mouse, taking 4 measurements from 5 different cross sections to obtain an average. Confocal images were taken on an Olympus FV3000 confocal microscope.

#### Glucose Tolerance Test

Mice were fasted for 4 hours and then injected with 100mg/mL D-glucose in sterile PBS (10uL/g body weight). Took blood glucose levels before injection, 15min, 30min, 45min, 60min, and 135min post-injection through the tail vein.

#### Mucus Thickness

7-μm thick slides were deparaffinized through a solution series for 5 minutes each (xylenes, xylenes, 100% EtOH, 95% EtOH, 75% EtOH, H_2_O). Alcian Blue/Pas staining was done using the protocol outlined in “Histotechnology: A Self-Instructional Text, 3^rd^ edition. Pages 115-116”. Slides were stained with Alcian Blue for 30 minutes, washed with running tap water for 2 minutes followed by a rinse with deionized (DI) water. Slides were stained with 0.5% Periodic acid for 5 minutes, washed with DI, then stained with Schiff’s Reagent for 30 minutes followed by a rinse with running tap water for 3 minutes. Nuclei were stained with Hematoxylin for 2 minutes, washed with running water for 30 seconds, followed by 10 dips in Clarifier, another 30 second wash, 10 dips in Bluing Reagent, and a final 30 second wash. Slides went through a solution series for 5 minutes each (95% EtOH, 100% EtOH, Xylenes, Xylenes). Coverslips were applied with Cytoseal and slides were allowed to dry overnight in the fume hood. The dense mucus layer of 40 locations evenly distributed from available cross sections on the slide (4-8 cross sections) were measured for each mouse at 60X magnification to obtain the average mucus thickness for each mouse, then combined to obtain the overall average and standard deviation.

#### Laser Capture Microdissection (LCM) and DNA extraction

Tissues were sectioned at 12μm and placed on Arcturus PEN Membrane Frame Slides and kept at 56°C overnight. Slides were deparaffinized through a solution series for 3 minutes each (xylenes, xylenes, 1:1; xylenes: EtOH, 100% EtOH, 100% EtOH, 95% EtOH, 70% EtOH, 50% EtOH, H_2_O). Slides were stained with Alcian Blue (1 min), rinsed, and Nuclear Fast Red (15 sec), then allowed to dry 2 hours. Regions of intestinal content were isolated and removed with the Applied Biosystems Arcturus Laser Capture Microdissection (LCM) system. Like-regions were pooled on LCM Caps from multiple cross sections of the same mouse until approximately 2,000,000 μm^2^ for 100-μm regions and 1,000,000 μm^2^ for 50-μm regions were collected. Isolated material was placed on Eppendorf tubes containing solutions from the QIAamp DNA Micro Kit with the addition of 3μL 10mg/mL lysozyme and incubated 16 hours at 56 °C. DNA was isolated following instructions of DNA Micro Kit and eluted with 30μL DNase-treated water.

#### Copy number of 16S rRNA per sample quantification

PCR amplified 16S rRNA gene from mouse fecal sample using Taq polymerase as described in the TOPO TA Cloning Kit User Guide with 16S rRNA Universal primers recognizing 340F and 514R (80). Cloned PCR product into a pCR 2.1-TOPO vector using the TOPO TA Cloning Kit and transformed recombinant vector into One Shot TOP10 Chemically Competent *E. coli* as described in the TOPO TA Cloning Kit User Guide. Isolated plasmid DNA with the Invitrogen PureLink Quick Plasmid Miniprep Kit. Checked 16S rRNA gene properly inserted into vector with PCR and EcoR1 restriction digest following protocol provided with restriction enzyme. Generated a standard curve using quantitative PCR and primers 340F and 515R with plasmid DNA of known concentrations, relating DNA concentration to copy number using the equation provided in Park and Crowley, 2005. Found the best fit line between CT value vs. copy number in order to relate the CT value of the unknown sample values to copy number.

#### 16S Amplicon Sequencing

Isolated DNA was amplified (30 cycles) using the Phusion High-Fidelity DNA Polymerase. Primers were designed to contain adapter overhangs complimentary for Nextera XT tagmentation with the locus-specific sequencing flanking the V4/V5 region of the 16S rRNA gene (518F and 926R). PCR products were cleaned with Ampure XP and visualized by agarose gel electrophoresis. A second round of PCR (up to 50ng of template DNA, 5-10 cycles) was performed to attach full indices and adapters using the Nextera XT Index Kit and Phusion HF Master Mix. PCR products from the second PCR were cleaned with Ampure XP and analyzed by agarose gel electrophoresis and the Agilent BioAnalyzer DNA 1000 chip. Quantification and normalization was performed on all samples prior to pooling using Qubit fluorometer and the final pooled library was quantified using qPCR in a Roche LightCycler 480 with the KAPA Biosystems Illumina Kit. Samples were sequenced (2×250 bp paired-end) on an Illumina MiSeq at the Rhode Island Genomics and Sequencing Center (Kingston, RI).

#### Bioinformatics

Quality control of raw sequences was performed using dada2 package v1.6.0 (81) in R v3.4.3. Forward and reverse reads were truncated to be 240bp and 190bp, respectively. The first 20bp were also removed to remove where primers bound constant regions, and standard filtering parameters were applied (82). Sequences were denoised by estimating error rates from pooled sequencing reads from all samples provided. Reads were then merged and chimeras identified and removed. Taxonomy was assigned using the RDP training set(83). The rooted phylogenetic tree was created using ape package v5.1 (84). Data were imported and analyzed using phyloseq v1.22.3 (85). Principal Coordinates of Analysis (PCoA) plots were made by calculating distances between communities using UniFrac (16) and transforming sample counts into percentages to normalize for unequal sampling depth. Alpha-diversity tests (Observed ASVs, Shannon, Pielou’s Evenness) were done on samples rarefied to the sample with the lowest sequencing depth. Raw values were exported to Prism to make final figures.

#### Quantitative PCR

Samples were prepared using Maxima SYBR Green/ROX qPCR Master Mix following the protocol outlined in the Maxima SYBR Green/ROX qPCR Master Mix User Guide. Relative abundance of Clostridia was determined using mix of CI-F1/CI-R2, CXI-F1/CXI-R2, and CIV-F1/CIV-R2 primer sets (86). Values were normalized to 16S rRNA Universal (340F and 515R).

#### Quantification and Statistical Analysis

Differences between groups of communities was calculated using a permutational multivariate analysis of variance (PERMANOVA) based on UniFrac distance (16). Differential abundance was calculated using DESeq2 (87), determining the adjusted p-value for multiple comparisons threshold as ≤ 0.05. Differences in blood glucose levels over time was tested with a 2-way ANOVA. All other significance tests were done using a parametric t-test, using a paired t-test when applicable.

## Data and Software Availability

Raw reads have been deposited in the NCBI Sequence Read Archive (SRA) database under the BioProject ID number PRJNA545452.

## Acknowledgments

We thank Dr. Namrata Iyer for helpful discussions, Dr. Geon Goo Han for assistance in conventionalization, Jenna Wurster for assistance with blood glucose level measurements, and Dr. Jim Mossman for statistical advice. We also thank Janet Atoyan for preparing libraries and sequencing, Geoffrey Williams for assistance with confocal microscopy, and members of the Brown University Histology Core, Paula Weston, Mindie Golde, and David Silverberg for sample preparation. This work was supported by NIH (P20GM10903 and 1R01DK113265 to S.V.) and DoD (W81XWH-18-1-0281 to S.V). S.V., K.D, and K. C. E designed experiments and performed analysis. S.V and K.D. wrote the manuscript.

---

**Supplementary Figure 1.**
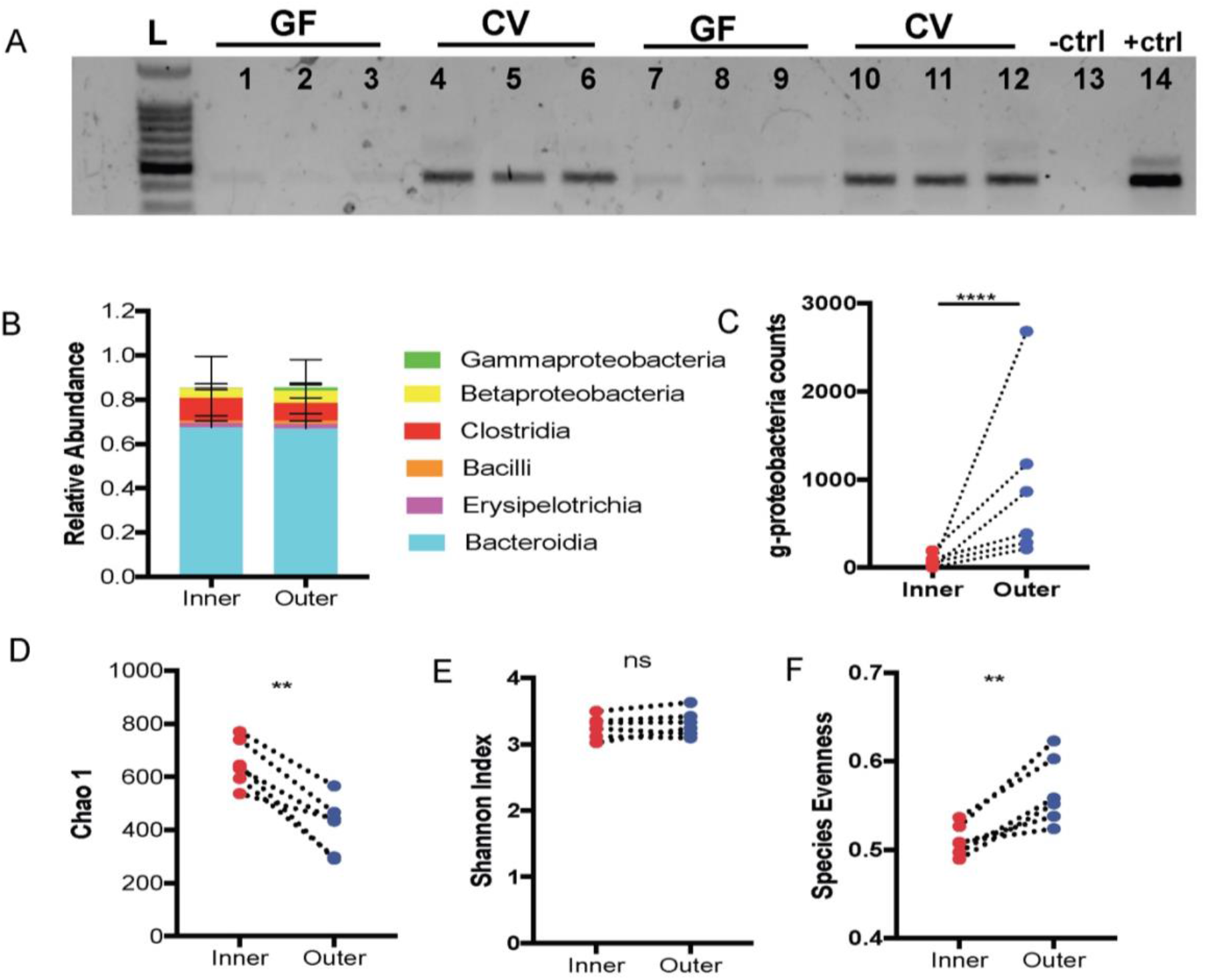
Characterization of dense community structure. A. Gel of PCR amplified samples extracted from 1^st^ 100-μm germ-free (GF) mice (lanes 1-3), 1^st^ 100-μm conventional mice (lanes 4-6), subsequent 50-μm GF mice (lanes 7-9), and subsequent 50-μm conventional mice (lanes 10-12) showing no PCR amplification from GF samples. Lane 13 negative control. Lane 14 positive control. B. Class level relative abundance of inner and outer communities (N = 6 mice from Cage 1). C. Differential abundance of class Gammaproteobacteria calculated using DESeq2, showing significantly more Gammaproteobacteria in the outer community (N = 6 mice from Cage 1) D. Chao1 diversity index showing higher richness in the inner community (N = 6 mice from Cage 1). E. Shannon diversity index showing no difference in Shannon diversity between inner and outer communities (N = 6 mice from Cage 1). F. Pielou’s Evenness index showing significantly higher evenness in the outer community (N =6 mice from Cage 1).

**Supplementary Figure 2.**
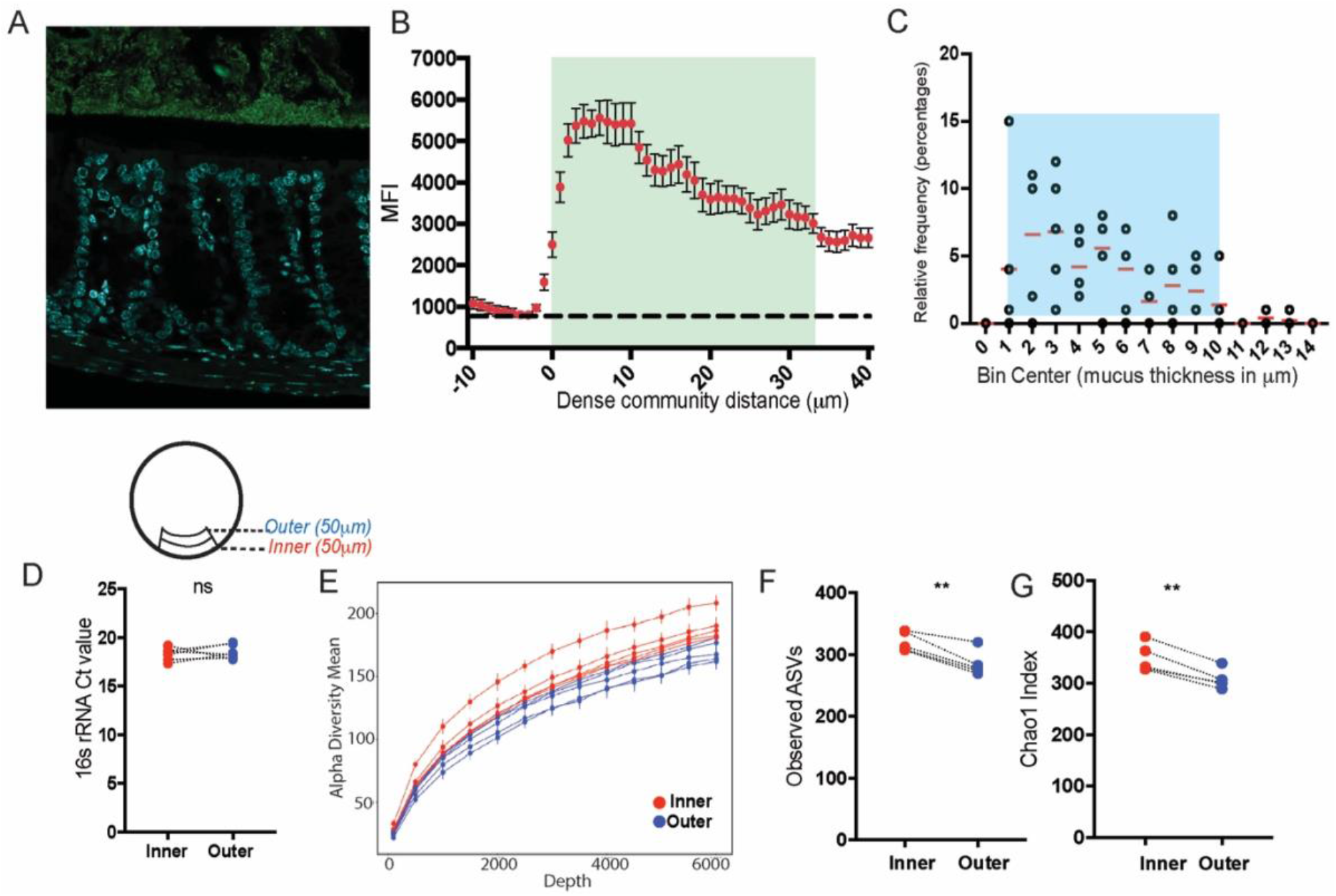
Dense community and mucus thickness in colon of Jackson mice. A. FISH image identifying all bacteria (green) to show biofilm-like structure close to host epithelium (DAPI, blue). B. Mean fluorescence intensity (MFI) measurements from a representative mouse to show bacterial density of biofilm-like structure (N = 3 mice). C. Relative frequency of dense mucus layer thickness measurements, described in Methods section under “Mucus Thickness” (N = 5 mice, 40 measurements each). D. Threshold cycle (Ct) values of 16s rRNA gene in inner and outer community. E. Alpha rarefaction curves plotting the number of unique amplicon sequence variants (ASVs) from inner (red) and outer (blue) communities showing more unique ASVs are found in the inner community (N = 5 mice). F. Observed ASVs showing significantly higher richness in the inner community (N = 5 mice). G. Chao 1 diversity index showing significantly higher richness in the inner community (N = 5 mice).

**Supplementary Figure 3.**
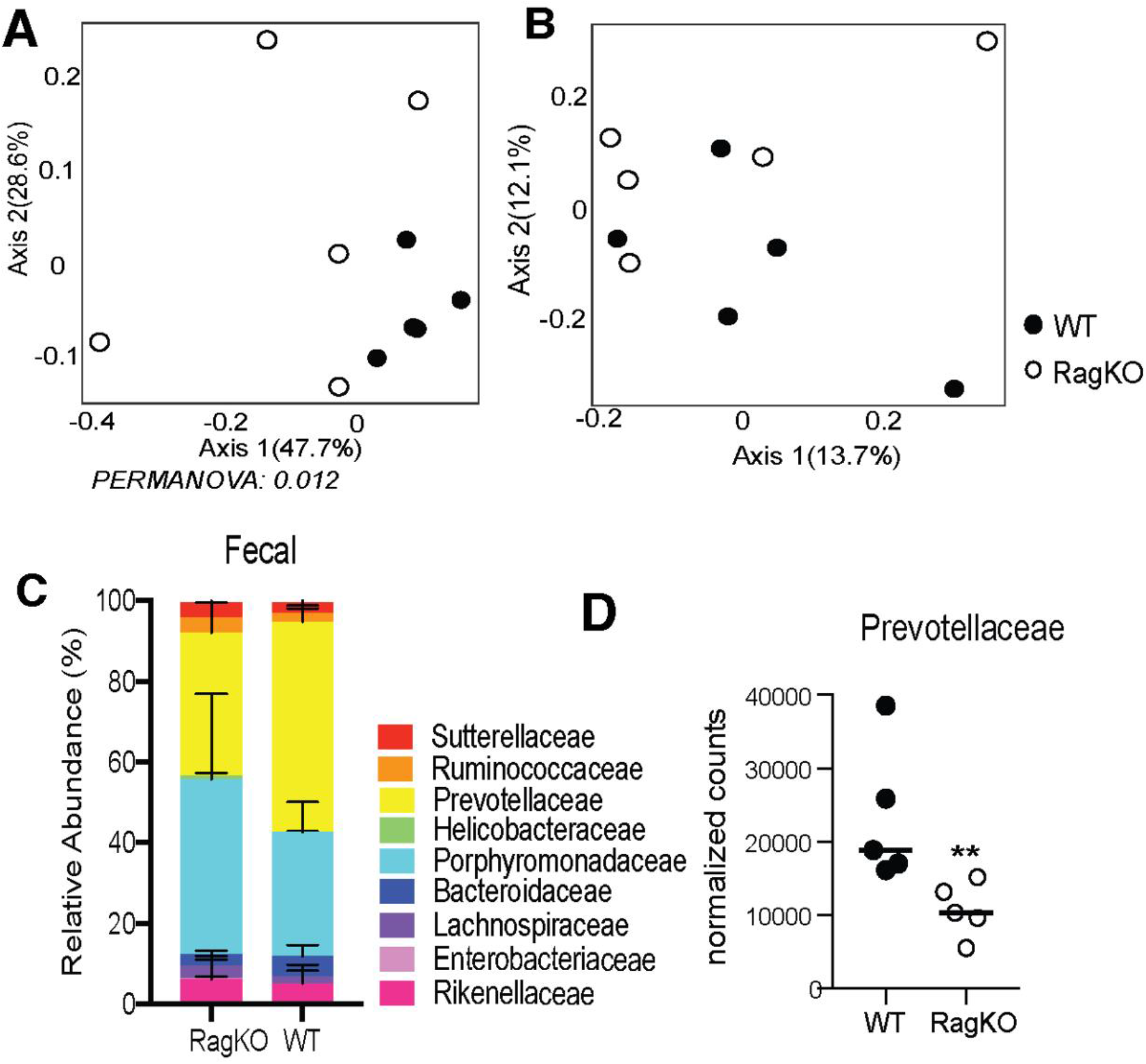
Differences in fecal communities between WT and Rag1KO mice. A. Principal coordinates of analysis (PCoA) plot using weighted UniFrac distances shows there is a significant difference between fecal communities of conventionalized WT and Rag1KO mice (N = 5 mice per group). B. Principal coordinates of analysis (PCoA) plot using unweighted UniFrac distances shows there is no significant difference between fecal communities of conventionalized WT and Rag1KO mice (N = 5 mice per group). C. Family level relative abundance of fecal communities in conventionalized Rag1KO and WT mice (N = 5 mice per group) D. Differential abundance of family Prevotellaceae calculated using DESeq2, showing significantly less Prevotellaceae in the fecal communities of Rag1KO mice (N = 5 mice per group)

**Supplementary Figure 4.**
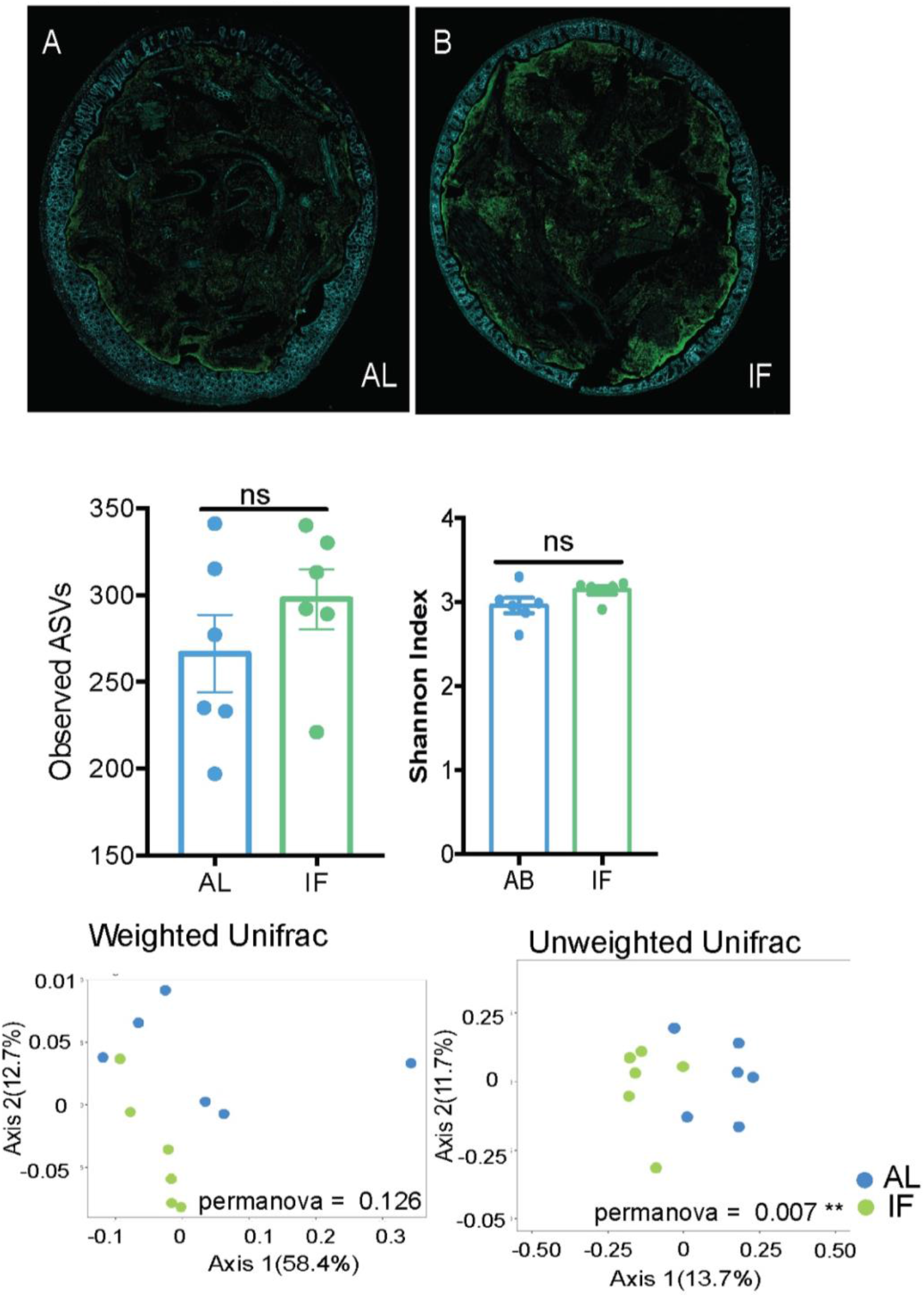
Comparison between AL and IF Fecal communities. A. Whole colonic cross section stained with fluorescent probes identifying all bacteria (green) and host epithelium (blue) of dense community structure in *Ad libitum* mice B. Whole colonic cross section stained with fluorescent probes identifying all bacteria (green) and host epithelium (blue) of dense community structure in Intermittent fasting mice. C. Observed ASVs found in AL and IF fecal communities showing no significant difference in richness between communities (N = 6 mice per group) D. Shannon diversity in AL and IF fecal communities showing no significant difference in diversity between communities (N = 6 mice per group) E. Principal coordinates of analysis (PCoA) plot using weighted UniFrac distances shows fecal communities from AL and IF are not significantly different (N = 6 mice per group) F. Principal coordinates of analysis (PCoA) plot using unweighted UniFrac distances shows fecal communities from AL and IF are significantly different (N = 6 mice per group).

